# HOX13-MEDIATED *DBX2* REGULATION IN LIMBS SUGGESTS INTER-TAD SHARING OF ENHANCERS

**DOI:** 10.1101/2020.11.16.379412

**Authors:** Leonardo Beccari, Gabriel Jaquier, Lucille Lopez-Delisle, Eddie Rodriguez-Carballo, Bénédicte Mascrez, Sandra Gitto, Joost Woltering, Denis Duboule

**Author notes:** Institut Neuromyogène. CNRS UMR 5310, INSERM U1217, University of Lyon, Lyon, France. Department of Molecular Biology, University of Geneva, 30 quai Ernest-Ansermet, 1211, Geneva, Switzerland (Eddie Rodriguez-Carballo). Zoology and Evolutionary Biology, Department of Biology, University of Konstanz, 78464, Konstanz, Germany (Joost Woltering). Corresponding authors. LB; DD /. **Conflict of interest disclosure:** The authors declare no competing interests.

## Abstract

**Background:** During tetrapod limb development, the HOXA13 and HOXD13 transcription factors are critical for the emergence and organization of the autopod, the most distal aspect where digits will develop. Since previous work had suggested that the *Dbx2* gene is a target of these factors, we set up to analyze in detail this potential regulatory interaction.

**Results:** We show that HOX13 proteins bind to eutherian-specific sequences at the vicinity of the *Dbx2* locus that have enhancer activity in developing digits. However, the functional inactivation of the DBX2 protein did not elicit any particular phenotype related to *Hox* genes inactivation in digits, suggesting either redundant or compensatory mechanisms. We report that the neighboring *Nell2* and *Ano6* genes are also expressed in distal limb buds and are, in part, controlled by the same *Dbx2* enhancers despite being localized into two different topologically associating domains (TADs) flanking the *Dbx2* locus.

**Conclusions:** We conclude that *Hoxa13* and *Hoxd* genes cooperatively activate *Dbx2* expression in developing digits through binding to eutherian specific regulators elements in the *Dbx2* neighborhood. Furthermore, these enhancers can overcome TAD boundaries in either direction to co-regulate a set of genes located in distinct chromatin domains.

**Bullet points:**

- *Hoxa13* and *Hoxd* genes cooperatively regulate *Dbx2* expression in developing digits via eutherian specific enhancers.
- *Dbx2* is expressed in different digit joint precursors but its function there is not essential.
- *Dbx2* enhancers also control the expression of the *Nell2* and *Ano6* genes, which are located in different TADs, thus overcoming the boundary effect.
- *Dbx2* chromatin architecture and enhancers evolved in the mammalian lineage.

**Grant Sponsor and Number:** Swiss National Research Foundation #310030B_138662.

European Research Council grants Regul*Hox* #588029

## INTRODUCTION

For many decades, the vertebrate limb has been an efficient experimental paradigm to study the basic principles and concepts underlying developmental processes. The main reason is the congruence between the definition of specific signaling regions in the developing limb buds, on the one hand, and their association with specific molecular markers, on the other hand. Classical experimental embryology indeed led to a fairly precise cellular definition of those regions in the limb bud, which have a particular activity and function, such as the limb apical ectodermal ridge and the zone of polarizing activity. John Fallon made seminal contributions in this early phase and was one of the pioneers of this field^1–3^; see also references in^4,5^, as well as^6–9^. Subsequently, transcription factors were cloned, which could be superimposed to such cellular landmarks, such as *Hox* genes ^10^ (see ^11^), followed by the key signaling molecules^12–14^. Soon after, gain- and loss-of-function experiments helped ascertain the central roles of these genes in controlling limb patterning and morphogenesis. In this view, the developing limb was the first vertebrate system where a bridge was established between cellular models and their molecular components.

Amongst these key factors are the *Hox* genes belonging to both the *HoxA* and *HoxD* clusters. They are transcribed into precisely delimited domains within the incipient limb buds ^10,15^ and they specify the proximal and distal limb segments as well as some anterior to posterior features ^16–18^; reviewed in ^19^. *Hoxa13* and *Hoxd13* are essentially required for the specification and development of the autopod, the distal-most limb domain that will give rise to the digits and part of the wrist. They control the size, shape, and number of autopod bones by regulating mesenchymal cell aggregation, chondrification and ossification^17,20–22^. In fact, the inactivation of both *Hoxa13* and *Hoxd13* in mice leads to an agenesis of the distal limb and the formation of a chondrogenic blastema at the distal extremity of the ulna/fibula and radius/ humerus ^17,18^. Different studies have addressed the identification of the HOXA13/HOXD13 downstream target genes in distal limb development. These included genes controlling cell adhesion, morphology, and proliferation/ survival (eg. *Hand2, Shh, EphA7, EphA3, Bmp2*/*Bmp7*; ^23–29^). The regulatory relationships and functional roles of many such target genes nevertheless remain poorly understood.

We previously reported that the transcription factor *Dbx2* (*Developing Brain Homeobox protein 2*) is strongly downregulated in distal forelimb cells lacking *Hox13* function ^23^. *Dbx2* belongs to the *Dbx* subfamily of homeobox-containing proteins and is expressed in the mouse embryonic brain and neural tube, as well as during limb development ^23,30,31^. However, while it is well established that *Dbx2* contributes to the specification of the V0 spinal cord interneurons ^32–35^, its potential role in limb development has remained elusive. However, a heterozygote deletion spanning the genomic region comprising the human *loci NELL2, DBX2* and *ANO6* was associated with intellectual retardation, skeletal and dental anomalies, reduced hand and feet size and clinodactyly of the fifth digit, suggesting that *Dbx2* could be involved in digit development ^36^, where it may mediate part of the functions of HOX13 proteins.

In this study, we characterized the regulation of the mouse *Dbx2* gene in developing digits. We show that the HOX13 factors directly regulate *Dbx2* expression in digits, in part by binding to eutherian-specific regulatory elements located within 30Kb 5’ to the *Dbx2* locus, as well as within its introns. Furthermore, we show that 5’ *Hoxd* genes also contribute to *Dbx2* regulation by acting cooperatively and redundantly with *HOX13* proteins. However, *Dbx2* null mice do not display any of the major limb skeletal abnormalities displayed by any combinations of *HOX* mutations, suggesting either that *Dbx2* is not a major downstream *Hox* effector or that its function is compensated for in this particular situation.

We also observed that the *Dbx2* neighboring genes *Nell2* (*Neural EGFL Like 2*) and *Ano6* (*Anoctamin 6*) are expressed in the distal limb as well. Analysis of chromatin interaction profiles revealed that at the 3D level, the *Nell2* and *Ano6* genes are organized into distinct Topologically Associating Domains (TADs), which are regions of the genome where gene-enhancer interactions are favored ^37^. Interestingly, the boundary between these two TADs maps at the proximity of the *Dbx2* locus and of its limb enhancers, which seem to be able to control the transcription of the three genes, regardless of in which TAD they reside.

## RESULTS

### *Dbx2* expression during distal limb development

We characterized *Dbx2* expression at different stages of mouse forelimb development by whole-mount in situ hybridization (WISH) and quantitative PCR (qPCR) and compared it with that of *Hoxa13* and *Hoxd13* (Fig. 1). Its transcripts were first scored during early limb development (E9.5-E10) throughout the limb bud mesenchyme with the exception of mesodermal cells underlying the distal-most limb ectoderm (Fig. 1A). This expression poorly correlates with that of *Hox13* genes and, overall, the *Dbx2* mRNA levels at this stage remained very low, as confirmed by qPCR analysis (Fig. 1B, C). By E11.5, *Dbx2* expression in the proximal limb was confined to a small and posterior moon-shaped domain of cells (Fig. 1A, asterisk). *Dbx2* transcripts were also detected in the anterior portion of the autopod (Fig. 1A, arrowhead). Thus, the distal limb expression of *Dbx2* was delayed by approximately 24h when compared to the onset of *Hoxa13* and *Hoxd13* in the autopod (Fig. 1A-C; ^10,38^). At E12.5, *Dbx2* mRNA spread to the entire distal-most portion of the autopod, both in digit and interdigit mesoderm and in a nested domain within the *Hoxa13*/*Hoxd13* expressing cells (Fig. 1A, B). However, *Dbx2* transcripts were rapidly downregulated in the interdigit region and, from E13 onwards, they were detected in sequentially formed domains reminiscent of the forming digit joints.

**Figure 1.**
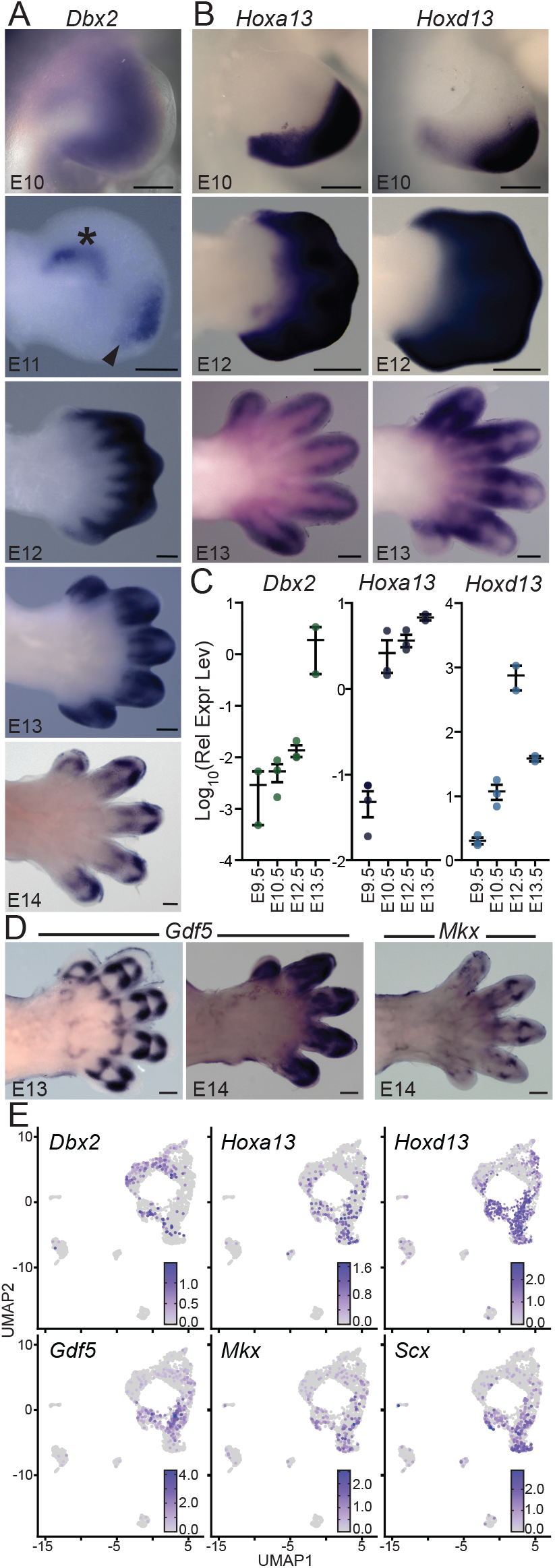
Analysis of *Dbx2* expression in developing digits. **A, B**. WISH analysis of *Dbx2* (A) and *Hoxa13*/*Hoxd13* (B) in mouse embryonic forelimbs at different developmental stages. Scale bar: 250µm. **C**. Quantitative PCR analysis of *Dbx2, Hoxa13* and *Hoxd13* mRNA levels in mouse forelimb buds, normalized to the housekeeping gene *Hmbs*. Each point represents independent biological replicates; bars represent the mean replicate value ± SEM. **D**. WISH analysis of *Gdf5* and *Scx* genes, which are expressed in cartilage and ligament precursors of the digit joint interzone domain and in tendon progenitors, respectively ^40,43^. Scale bar: 250µm. **E**. UMAP representation of the scRNA-seq data from mouse E13 mouse hindlimbs ^39^, showing the expression of *Dbx2, Hoxa13* and *Hoxd13*, as well as of different joint (*Gdf5*) and tendons/ligaments (*Mkx, Scx*) markers ^40,41,43^.

To assess which cell type(s) express the *Dbx2* gene, we used known markers of tendon and cartilage precursors expressed in digit joints. We also re-analyzed single cell-RNA sequencing (scRNA-seq) datasets from E11, E13 and E15 mouse hindlimbs ^39^ (Fig. 1D, E; Fig. S1). This analysis revealed that *Dbx2* is expressed in different cell populations within the developing limb. Some positive cells did not express *Hoxa13*/ *Hoxd13* at the stages analyzed and displayed the *Col2a1* marker of mature cartilage precursors (Fig. 1D; Fig. S1A). *Dbx2* transcripts were also detected in a subpopulation of *Hoxa13* and *Hoxd13* positive cells, which expressed the *Gdf5, Mkx, Scx and Col1a1* genes as well (Fig. 1D, E; Fig. S1A). *Gdf5* is transcribed in different cartilage and tendon-ligament precursors of the joint interzone, whereas *Mkx, Scx* and *Co1a1* mark tendon cell precursors ^39–43^. Of note, although *Dbx2/Col2a1/Sox9+* cartilage cells did not express *Hox13* genes at E13/E15, they derive from a common population of precursors expressing *Hox13* genes (Fig. S1B)^39^.

These results showed that *Dbx2* is expressed during digit development in *Hoxa13*/ *Hoxd13* expressing cells corresponding to tendon and cartilage precursors of developing digit joints. Therefore, it supports the possibility that HOX13 proteins could act as direct regulators of *Dbx2* expression in these cells, in agreement with the reported function of 5’*Hoxd* and *Hoxa13* genes in digit joint development ^40,44,45^.

### Identification of HOX13-bound sequences regulating *Dbx2* expression in digits

*Dbx2* expression in distal limbs is strongly compromised in the absence of HOX13 proteins ^23^. To further evaluate whether HOX13 paralogs could act as direct regulators of *Dbx2* expression, we set up to characterize the extent of the *Dbx2* regulatory landscape both by analyzing available Hi-C datasets and by performing 4C-seq experiments (Fig. 2). TADs are megabase-scale structures that constitute a unit of 3D genome organization ^37,46^. Thus, they often coincide with and delimit the extent of gene regulatory landscapes ^47,48^. TADs are generally independent from the transcriptional status of the gene(s) inside and can be identified across different cell types or tissues.

**Figure 2.**
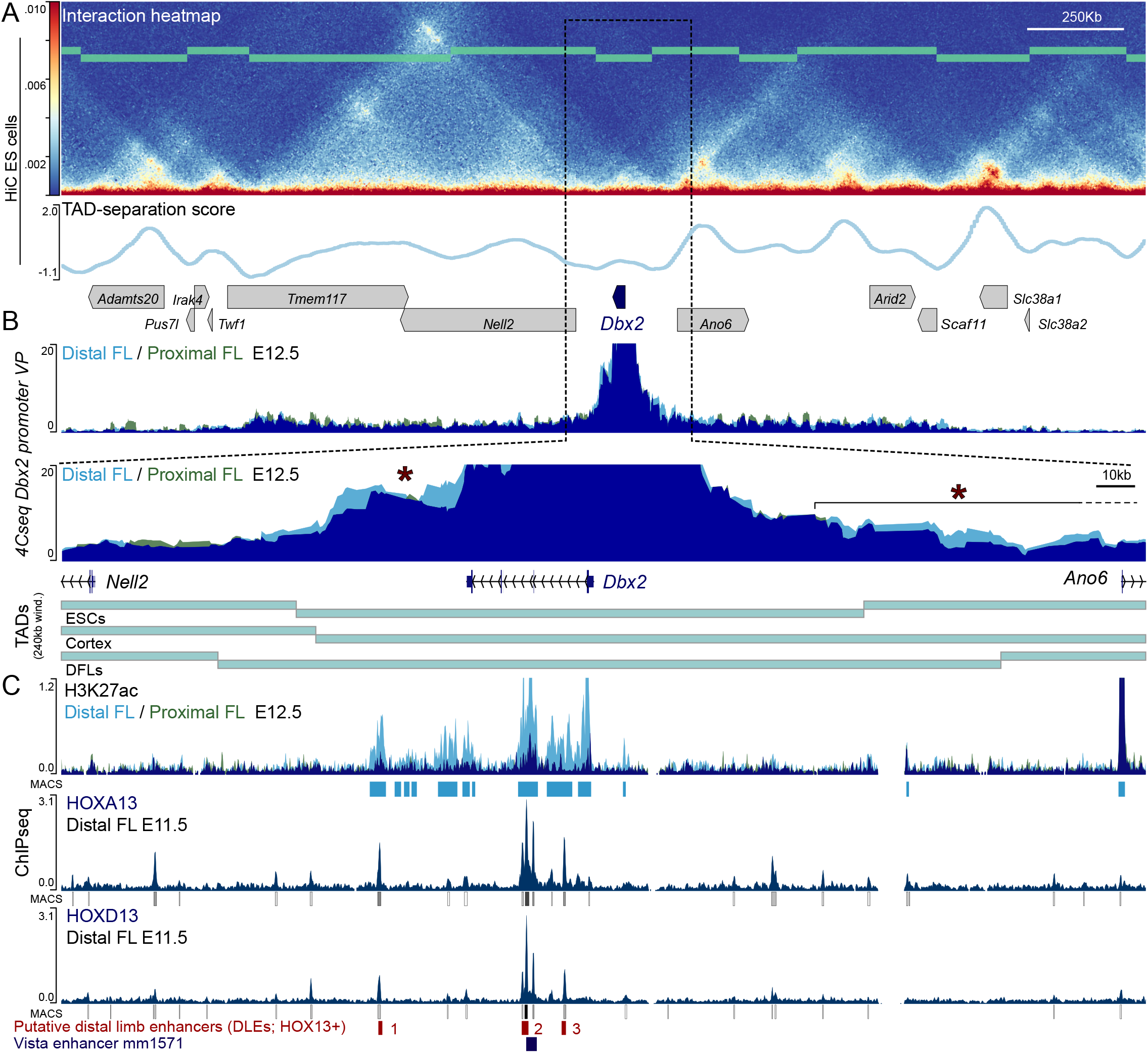
The *Dbx2* regulatory landscape in mouse limb buds. **A**. High resolution (5Kb bin size) Hi-C map (top) of the *Dbx2* genomic region in mouse ES cells and graphs showing the TAD-separation score (bottom); windows size 120Kb. Protein coding loci are represented by blue (*Dbx2*) or gray boxes (all other genes) pointing towards the gene direction. Data from ^49^. **B**. 4C-seq analysis of *Dbx2* interactions using the *Dbx2* promoter as a viewpoint in E12 mouse distal (light blue) and proximal (green) forelimbs. Profile overlap is in dark blue. Each curve marks the average profile of three independent biological replicates. Asterisks mark the region(s) displaying increased contact frequencies in the DFL *versus* PFL. TADs in A and B are depicted with thick green lines (see also Fig. S2). **C**. ChIP-seq analysis of H3K27ac marks in distal (light blue) and proximal (green) forelimbs, and HOXA13/HOXD13 binding profiles over the *Dbx2* genomic region (below) and their respective peak calling. Profile overlap is in dark blue. Data from ^23,28^. HOX13+ putative regulatory elements (DLE1 to 3) are shown in red. The Vista enhancer mm1571 is represented by a blue rectangle.

We analyzed high-resolution (5Kb bin size) Hi-C data from ES cells and embryonic cortex ^49^, as well as 40Kb-resolution Hi-C profiles from E12 mouse distal limbs ^50^ (Fig. 2A; Fig. S2). We observed that, in all cases, the *Dbx2* genomic region is organized in two large TADs, which span the neighboring loci *Tmem117* and *Nell2* (5’TAD) and *Ano6, Arid2* and *Scaf11* (3’TAD), respectively. Although with some variation between tissues or cell types and depending on the TAD-separation score calculation parameters, the border between these two TADs consistently falls in the close vicinity of the *Dbx2* gene. Accordingly, a region of approximately 150Kb spanning the *Dbx2* locus forms a micro-domain of higher contact frequency spanning the TAD boundary (hereafter referred to as interTAD domain) and the *Dbx2* interactions were mostly restricted to this interTAD domain.

To confirm this, we performed 4C-seq experiments in mouse proximal and distal forelimb cells at E12 using the *Dbx2* promoter as a viewpoint (Fig. 2B). As expected, the vast majority of *Dbx2* interactions were observed within the 150Kb region, matching the interTAD domain identified in the Hi-C data analysis. However, some diffuse *Dbx2* interactions were also detected over the entire lengths of the 5’ and 3’ TADs flanking the *Dbx2* locus, while *Dbx2* contacts dramatically dropped down outside of these domains. Overall, the *Dbx2* interaction profiles remained very similar in both proximal and distal forelimbs (PFL and DFL, respectively). Nonetheless, we scored a DFL-specific increase in contacts over a narrow region located 55Kb away from the *Dbx2* transcription start site (TSS) on the 5’ side of the locus, as well as with a broad region comprised between 82Kb and 236Kb 3’ to the *Dbx2* transcription start site located and encompassing part of the neighboring *Ano6* gene (Fig. 2B, asterisk and Fig S3A). These results suggested that *Dbx2* expression in the developing limbs is mostly driven by mid and short-range regulatory interactions within its immediate 150Kb surroundings.

To identify putative regulatory sequences controlling *Dbx2* expression in developing digits, we analyzed H3K27 acetylation datasets ^23^, a histone modification specifically enriched in active enhancers and promoters ^51^. In mouse E12 PFL and DFL cells, within the 800Kb spanned by *Dbx2* and its flanking TADs, we identified only five non-coding regions specifically enriched in H3K27ac (Fig. 2C and Fig. S3B). These were located within the *Dbx2*-interTAD domain, suggesting that they could correspond to *Dbx2* regulatory elements. Of these, two were located in the intergenic region on the 5’ side of the *Dbx2* locus, two others mapped within *Dbx2* intronic sequences and one overlapped with the first *Dbx2* exon and TSS. All these sequences were strongly contacted by the *Dbx2* promoter (Fig. 2B). Other H3K27ac-positive regions were identified within the *Ano6*/*Arid2*/*Scaf11* TAD, yet they were not specifically enriched in this epigenetic mark in DFL cells, arguing against a specific involvement of these sequences in *Dbx2* regulation.

We also analyzed HOXA13 and HOXD13 ChIP seq datasets ^28^ to determine whether these proteins would directly interact with the *Dbx2* locus (Fig. 2C and Fig. S3). We observed HOXA13/HOXD13 binding at several locations within the *Dbx2* genomic region. While most of these HOX13 bound sequences were not located in H3K27ac-positive and *Dbx2* interacting regions, we nevertheless observed strong binding of these proteins in three of the DFL-specific H3K27ac-positive regions showing an interaction with *Dbx2*. One of these HOX13-bound sequences partially overlapped with a Vista enhancer (mm1571) previously characterized to drive *LacZ* reporter expression in the neural tube and developing limbs ^52,53^ (arrowhead in Fig. 3B). We quoted the other sequences as putative Distal Limb Enhancers (DLE) and numbered them based on their 5’to 3’ position within the *Dbx2* interacting domain (DLE1 to DLE3). These sequences are conserved across the different mammalian species analyzed, with the exception of DLE1, which is absent from the *Dbx2* genomic region in ungulate species, suggesting a specific loss of this element in this taxon (Fig. 3A). Instead, the Vista mm1571 enhancer was conserved in all tetrapod species analyzed. Furthermore, we could identify evolutionarily conserved HOX binding sites within the DLE1 and DLE2 sequences (Fig. 3A).

**Figure 3.**
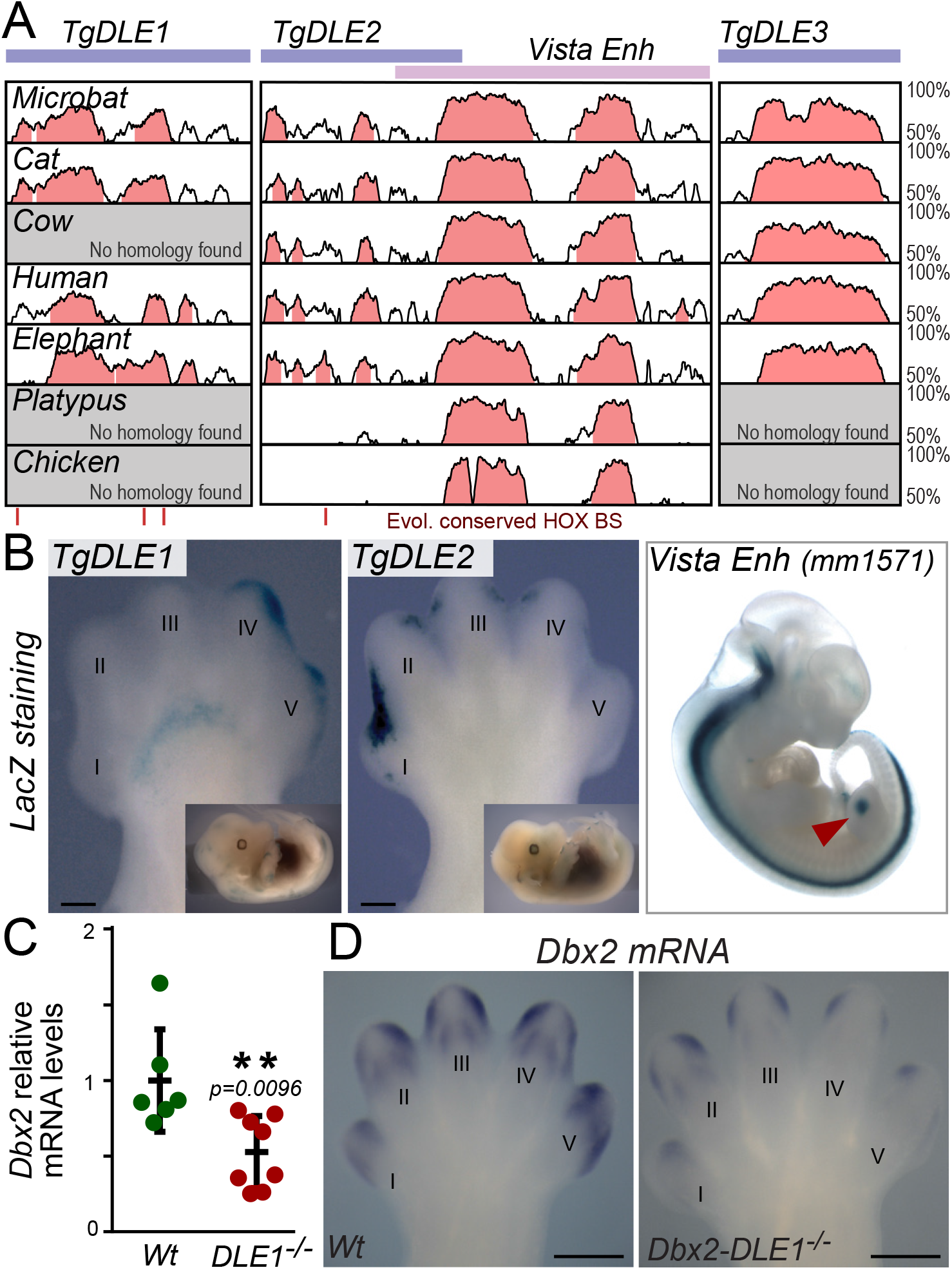
Putative *Dbx2* enhancers are active in distal limb buds. **A**. Vista alignment of the DLE1-3 regions and of the previously reported mm1571 regulatory element (depicted by purple and pink boxes, respectively). Evolutionarily conserved HOX binding sites are marked by vertical red lines at the bottom. **B**. X-gal staining of embryo transgenic for the DLE1 and DLE2 regulatory sequences in E13 mouse forelimbs (left) and of the mm1571 Vista enhancer (image from Vista enhancer browser; https://enhancer.lbl.gov/ ^53^). Distal limb expression of the mm1571 enhancers is marked by a red arrowhead. **C, D**. Quantitative PCR (C) and WISH (D) analysis of *Dbx2* expression in the distal forelimb of wildtype (Wt) and DLE1^-/-^ littermates. Each point represents independent biological replicates; bars represent the mean ± SEM. Values are normalized to the *Hmbs* gene and to the wt. In B and D, Digits are numbered (I-V) in the antero-posterior order. Scale bar: 250µm.

To assess the functional role of these DLEs, we cloned the DLE1 and DLE2 sequences in a *LacZ* reporter vector and tested them in transient transgenic experiments. The two elements displayed activity in E13 DFLs in a domain reminiscent of *Dbx2* expression in the last forming joint of the phalanges (Fig. 3B). Interestingly, DLE1 and DLE2 displayed mirror-imaged stainings, with DLE1 active in the posterior portion of the handplate and DLE2 anteriorly. Besides, DLE1 displayed weak yet reproducible activity in a narrow stripe of cells within the mesopod (Fig. 3B, asterisk), possibly related to the initial expression of *Dbx2* at E11.5 (Fig. 1A, asterisk). Likely, this was maintained until E13 due to the stability of the beta-galactosidase protein. Neither DLE1 nor DLE2 displayed transgene activity in any embryonic structure other than the developing digits. To corroborate the functional role of the identified elements on *Dbx2* regulation, we used CRISPR/Cas9 genome editing to produce mice lacking the DLE1 regulatory element (DLE1^-/-^) and analyzed *Dbx2* expression. As expected, DLE1^-/-^ mice displayed a significant decrease in *Dbx2* expression in the E13 developing digits, as compared to control littermates (Fig. 3C,D). In agreement with the DLE1 transgenesis results, this effect was even more pronounced in the posterior digits, where the DLE1 transgene displayed *LacZ* activity.

### *Hoxa13* and 5’ *Hoxd* genes directly regulate *Dbx2* expression

To assess the relative contribution of *Hox13* paralogs to *Dbx2* regulation, we measured its expression in the forelimb autopods of either *Hoxa13*^-/-^ or *Hoxd13*^-/-^ mice, and of compound mutants carrying different combinations of *Hoxa13* and *Hoxd13* null alleles (Fig. 4A), by using both WISH and qPCR. As previously described, *Dbx2* was almost completely abrogated in double *Hox13* mutant mice (Fig. 4B, C). Instead, only a weak reduction in *Dbx2* expression was observed in *Hoxd13*^-/-^ single mutants. There, transcripts were maintained in the distal forelimb, with the exception of the posterior-most portion of the autopod, where *Dbx2* expression was sharply reduced (Fig. 4B, C). In contrast, *Dbx2* mRNA levels strongly decreased in *Hoxa13*^-/-^ embryos and transcripts remained detectable at low levels only in the distal portion of the central digits. *Dbx2* expression was not detected in either the *Hoxa13*^-/-^*Hoxd13*^+/-^ or *Hoxa13*^+/-^*Hoxd13*^-/-^ compound mutants (Fig. 4B), indicating that a single allele of either genes was not sufficient to activate *Dbx2* in the DFL, despite the fact that in these mutants, a reduced but correctly specified autopod is still observed ^17,23^. Of note, a very faint and spatially ill-defined *Dbx2* signal was scored in the *Hox13* double knock-out mice (Fig. 4B), reminiscent of the early expression of *Dbx2* in the incipient limb bud at E9.5 to E10 (Fig. 1A). This expression was not observed in either *Hoxa13*^-/-^*Hoxd13*^+/-^ or *Hoxa13*^+/-^*Hoxd13*^*-/-*^ mutant embryos. This may reflect the inability of *Hox13* mutant limbs to properly terminate the early limb developmental program and to initiate the transcriptional network operating at later stages in the autopodial domain ^23,28^.

**Figure 4.**
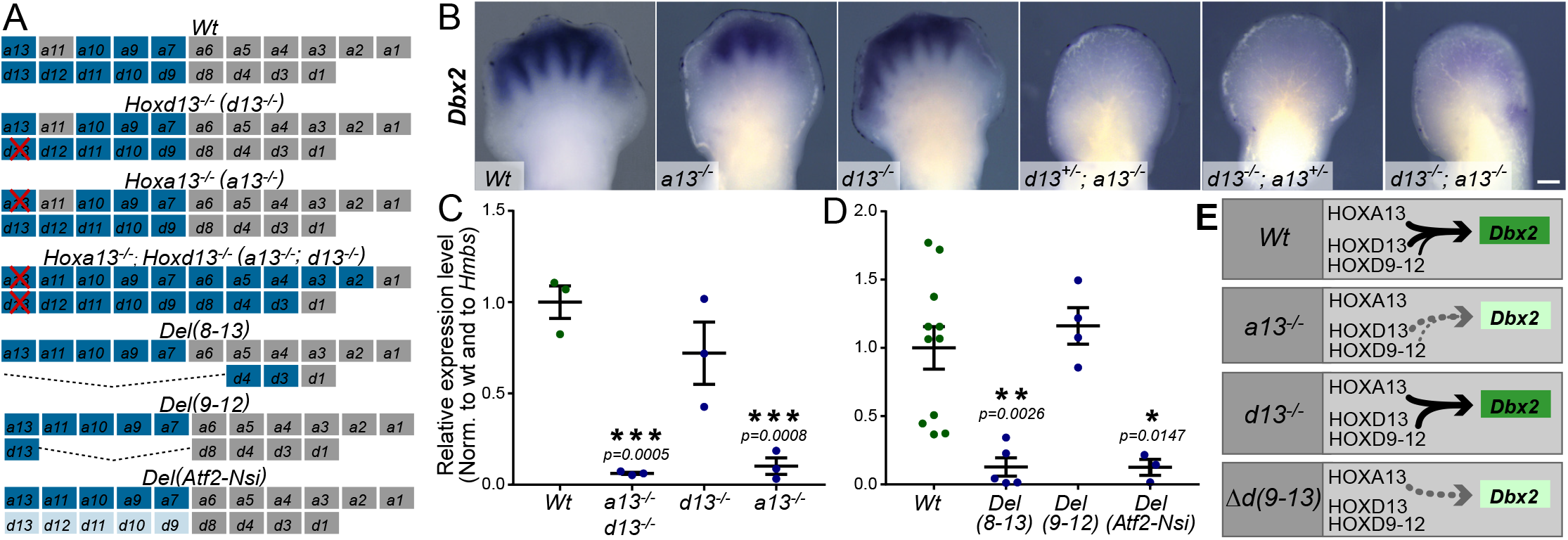
HOXA13 and 5’ HOXD proteins cooperatively regulate *Dbx2* expression in developing digits. **A**. Scheme of the different *Hoxa* and *Hoxd* paralogs expressed (in blue) in Wt distal limbs and in those of mice carrying homozygote mutations disrupting or altering the expression of the mouse *Hoxa13* and *Hoxd* paralogs. Silent genes are in gray. Red crosses represent inactivated genes. Dashed lines represent various deletions at the *HoxD* locus. The Del(*Atf2-Nsi*) mice carry a large genomic deletion spanning the centromeric TAD flanking *HoxD*. They display virtually no expression of any *Hoxd* genes in digits. **B**. WISH analysis of *Dbx2* expression in E12 mouse forelimbs of control and compound *Hoxa13*/*Hoxd13* mutant mice. Scale bar: 250µm. **C**,**D**. Quantitative PCR analysis of *Dbx2* expression in the DFL of control embryos or in different *Hoxa13, Hoxd13* and *HoxD* mutant alleles. Each point represents independent biological replicates; bars represent the mean replicate value ± SEM. P-values are calculated based on t-test comparison against Wt values **E**. Model explaining the cooperative role of *Hoxa13* and *Hoxd* genes in *Dbx2* regulation. Arrow thickness represents the relative contribution of each HOX protein. Gray dashed arrows depict weak *Dbx2* activation.

Because *Hoxd* genes exert largely overlapping functions in the development of the distal limb domain ^54^, we also assessed whether other *Hoxd* paralogs could contribute to *Dbx2* regulation. We thus analyzed *Dbx2* expression in series of mutant mice carrying deletions of different combinations of *Hoxd* genes (Fig. 4A, D). *HoxD*^*Del(Hoxd9-Hoxd12*)^ knock-out mice, hereafter referred to as Del(9-12), carry a deletion including all *Hoxd* genes normally expressed in the autopod but *Hoxd13*. They displayed normal levels of *Dbx2* expression as compared either to control or to *Hoxd13*^*-/-*^ mutant autopods. Instead, mice carrying a homozygote deletion including from *Hoxd8* to *Hoxd13 (HoxD*^*Del(Hoxd8-Hoxd13*)-/-^; hereafter Del(8-13)) displayed a drastic downregulation of *Dbx2* mRNA levels, which was significantly stronger than that observed in *Hoxd13*^*-/-*^ mutant and comparable to that of *Hoxa13*^*-/-*^ mice. This reduction was also observed in mice carrying a large genomic deletion removing the *HoxD* centromeric gene desert, which contains all the elements controlling *Hoxd* gene expression in the autopod ^55^. Altogether, these data indicate that although *Hoxd13* is the main *Hoxd* gene regulating *Dbx2* expression in digits, other *Hoxd* genes cooperatively contribute to this activation along with *Hoxa13* (Fig. 4E).

### *Dbx2* does not significantly contribute to the distal limb skeleton development

To determine the extent to which *Dbx2* contributes to HOX13 functions in distal limbs, and also to assess its importance in the hand/foot phenotype associated with the deletion of the human *NELL2/DBX2/ANO6* genomic region ^36^, we used CRISPR/Cas9 genome editing to disrupt the *Dbx2* homeodomain (Fig. 5A). We designed specific sgRNAs targeting the flanking region of the *Dbx2* third exon, which encodes two out of the three alpha-helices (H1-H2) of the homeodomain and part of the third (H3). We thus produced mice carrying a 377 bp large deletion, which removes the H1-2 coding sequence and produces a frameshift mutation, thus disrupting also H3 and the DBX2 C-terminal domain. This mutation is expected to prevent the binding of the protein to DNA and hence inactivate its function (Fig. 5A, B). The frequency of mouse pups carrying this *Dbx2* mutant allele, either heterozygous or homozygous, was significantly reduced when compared to the expected Mendelian ratio (Fig. 5C), suggesting that the *Dbx2* mutation led to embryonic or perinatal lethality. However, no clear skeletal or hand/ foot phenotype was observed in the *Dbx2*^*-/-*^ mice, neither in the length or number of phalanges, nor in their degree of ossification or in their phalangeal joints (Fig. 5D, F). Therefore, although *Dbx2* operates downstream of HOX13 genes in distal limb development, it is not the main contributor to the effects observed in these structures upon the loss of *Hox13* and other *Hoxd* genes ^17,40,45^.

**Figure 5.**
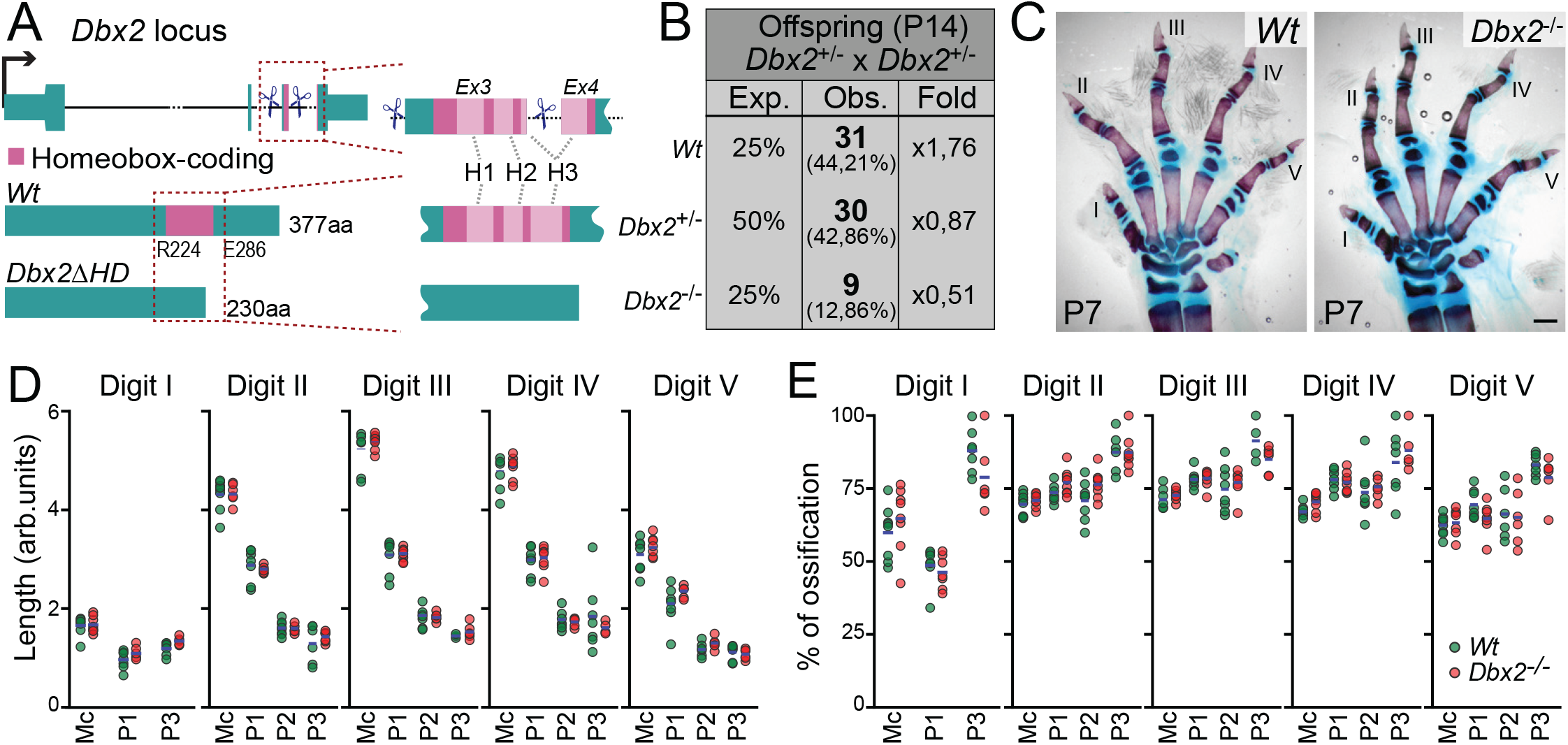
Disruption of the DBX2 homeodomain. **A**. *Dbx2* locus structure and predicted proteins in control and mutated *Dbx2* alleles. Blue scissors indicate the sgRNAs used for CRISPR/Cas9 genome editing. The homeodomain-coding portion is highlighted in pink. The first and last aminoacid positions of the DBX2 homeodomain are indicated. Its three *α*-helices (H1-3) are depicted by light pink boxes. Gray lines represent the correspondence between the DBX2 H1-H3 and its encoding sequence at the *Dbx2* locus. **B**. Table showing the proportion of *Dbx2*^+/+^, *Dbx2*^+/-^ and *Dbx2*^-/-^ P14 offspring obtained from *Dbx2*^+/-^ X *Dbx2*^+/-^ crosses. **C**. Alcian blue and alizarin red staining of the forelimb of P7 Wt or *Dbx2*^-/-^ littermates. Digits are numbered (I-V) in the antero-posterior order. Scale bar: 500µm. **D, E**. Quantification of the length (D) and degree of ossification (E) of metacarpals (Mc) and phalanges (P1-P3) of digits I-V in Wt (green) or *Dbx2*^-/-^ (red) littermates. Bone length was calculated as the distance between the tips of the epiphysis. The degree of ossification was calculated as the ratio of the length of the alizarin red + domain and the total bone length. Each point represents a biological replicate. Blue lines depict the mean of all biological replicates.

### *Nell2*/*Dbx2*/*Ano6* coregulation in developing limbs

The absence of limb alterations in *Dbx2* null mice raised the question of whether the neighboring *Nell2* and *Ano6* genes may contribute to limb development. In fact, the entire *Dbx2* genomic region has a syntenic interval in humans and other tetrapods (Fig. 6A) and the deletion involved in hand-foot defects in humans also contains the *NELL2* and *ANO6* genes ^36^. WISH analysis as well as mining a scRNA-seq dataset ^39^ revealed that *Nell2* and *Ano6* are specifically expressed in the distal portion of mouse developing limbs, in a population of *Hoxa13*/*Hoxd13* double-positive cells, part of which also express *Dbx2* (Fig. 6B, C). In both cases, their transcripts were distributed on both sides of the developing digits, displaying an indentation (*Nell2*) or a faint band (*Ano6*) corresponding to the joints of the forming phalanges (Fig. 6B and Fig. S3B). However, we could not identify any DFL-specific H3K27ac positive region in the *Nell2* or *Ano6* TADs (Fig. 2B; Fig. S3A, B), suggesting that their expression in developing limbs could be driven by the regulatory elements of the *Dbx2*-containing interTAD region.

**Figure 6.**
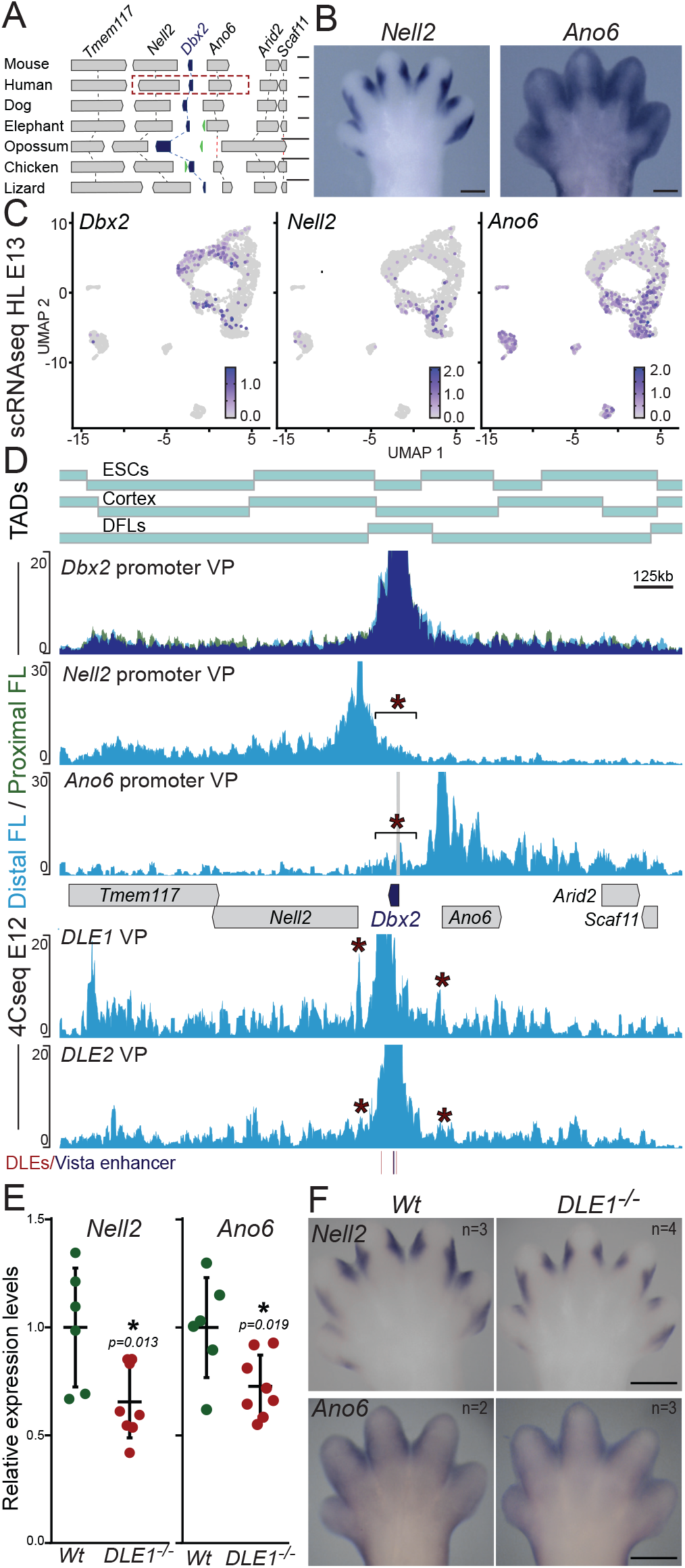
Expression and regulation of *Nell2* and *Ano6* in mouse limb buds. **A**. Synteny of the *Dbx2* genomic region. *Dbx2* is in blue and other genes in gray boxes. The red dashed rectangle depicts the deleted region reported in a human case ^36^. Gray and red dashed lines indicate syntenic relationship and orthologous gene loss, respectively. Gene insertions are in green. Scale bar: 100Kb. **B**. WISH of *Nell2* and *Ano6* in E13 mouse forelimbs. **C**. UMAP representation of the scRNA-seq data from E13 mouse hindlimbs ^39^ for *Dbx2, Nell2* and *Ano6*. **D**. 4C-seq profiles showing the interactions of *Dbx2* (see also Fig. 2A), *Nell2, Ano6*, DLE1 and DLE2 in proximal (green) and/or distal (light blue) forelimbs (profile overlap is in dark blue). The gray contacts in the *Ano6* viewpoint correspond to probably artefactual PCR product. *Nell2, Ano6*, DLE1 and DLE2 profiles are from a single experiment. **E, F**. Quantitative PCR (**E**) and WISH analysis (**F**) of *Nell2* and *Ano6* expression in E13 DFL of control and DLE1^-/-^ embryos. Each point represents independent biological replicates; bars represent the mean ± SEM. Values are normalized to the *Hmbs* gene and to the wt. Scale bar in B and F: 250µm.

To address this question, we performed 4C-seq experiment in E12 DFLs using either the *Nell2* or the *Ano6* promoters, as well as DLE1 and DLE2 as viewpoints (Fig. 6D; Fig. S3D, E). As expected, the *Nell2* and *Ano6* promoters displayed strong interactions with sequences located in their own TADs, while they showed reduced contacts with the neighboring TAD. However, in both cases, we observed significant contacts of both the *Nell2* and *Ano6* promoters with the DLE1-3 region (Fig. 6D and Fig. S3C). In the reverse experiment, DLE1 interacted not only with the *Dbx2* promoter, but also with the close neighborhood of the *Nell2* and *Ano6* TSSs. Such interactions were also scored when using DLE2 as a viewpoint, though with reduced in intensity, likely due to the fact that the DLE2 contacts remained overall strongly confined to the *Dbx2* interTAD domain. Nonetheless, DL2 contacts with *Nell2*/*Ano6* were still higher than those displayed by the *Dbx2* promoter, which contacted predominantly the interTAD region (Figs. 2B, 6D and Fig. S3C). We did not observe any significant enrichment of H3K27me3, a histone modification usually associated with inactive enhancers and promoters ^56^, over the DLE1 to 3 regions, ruling out the possibility that the interactions would represent contacts between H3K27me3 islands, as reported in other instances ^57^.

To further document that part of the transcription of both *Nell2* and *Ano6* could be driven by elements shared with the *Dbx2*, we analyzed their expression in mice lacking the DLE1 sequence by WISH and qPCR. We observed that *Nell2* and *Ano6* transcript levels were significantly decreased in the autopods of DLE*1*^*-/-*^ embryos when compared to control littermates (Fig. 6E, F). This decrease was more pronounced for *Nell2* than for *Ano6*, yet it remained proportionally lower than that observed for *Dbx2* (Fig. 3C, D), in agreement with the differences observed in contact frequency between DLE1-2 and the promoters of these three genes. These results strengthened the hypothesis that the regulatory elements located in the genomic vicinities of *Dbx2* also control part of the *Nell2* and *Ano6* expression in developing digits.

### Structural differences at the *Nell2*/*Dbx2*/*Ano6* locus between birds and eutherian mammals

While the DLE1-3 regulatory elements are broadly conserved across eutherians, they could neither be identified in non-eutherian mammals, nor in any other vertebrate (Fig. 3A). Instead, a large syntenic region around the *Dbx2* locus is conserved across all tetrapods, except in monotremes, where the *Ano6* gene was specifically lost (Fig. 6A). Hi-C interaction profiles produced from embryonic chicken limbs ^58^ revealed that the TAD organization of the chicken *Dbx2* region is similar to that of the mouse (Fig. 2A, 7A and Fig. S2), with *Dbx2* located in the close vicinity of the boundary between the *Nell2* and *Ano6* containing TADs (Fig. 7A), in agreement with the syntenic correspondence. This suggests that the *Dbx2* TAD architecture is maintained across vertebrates and arised before the emergence of mammals.

**Figure 7.**
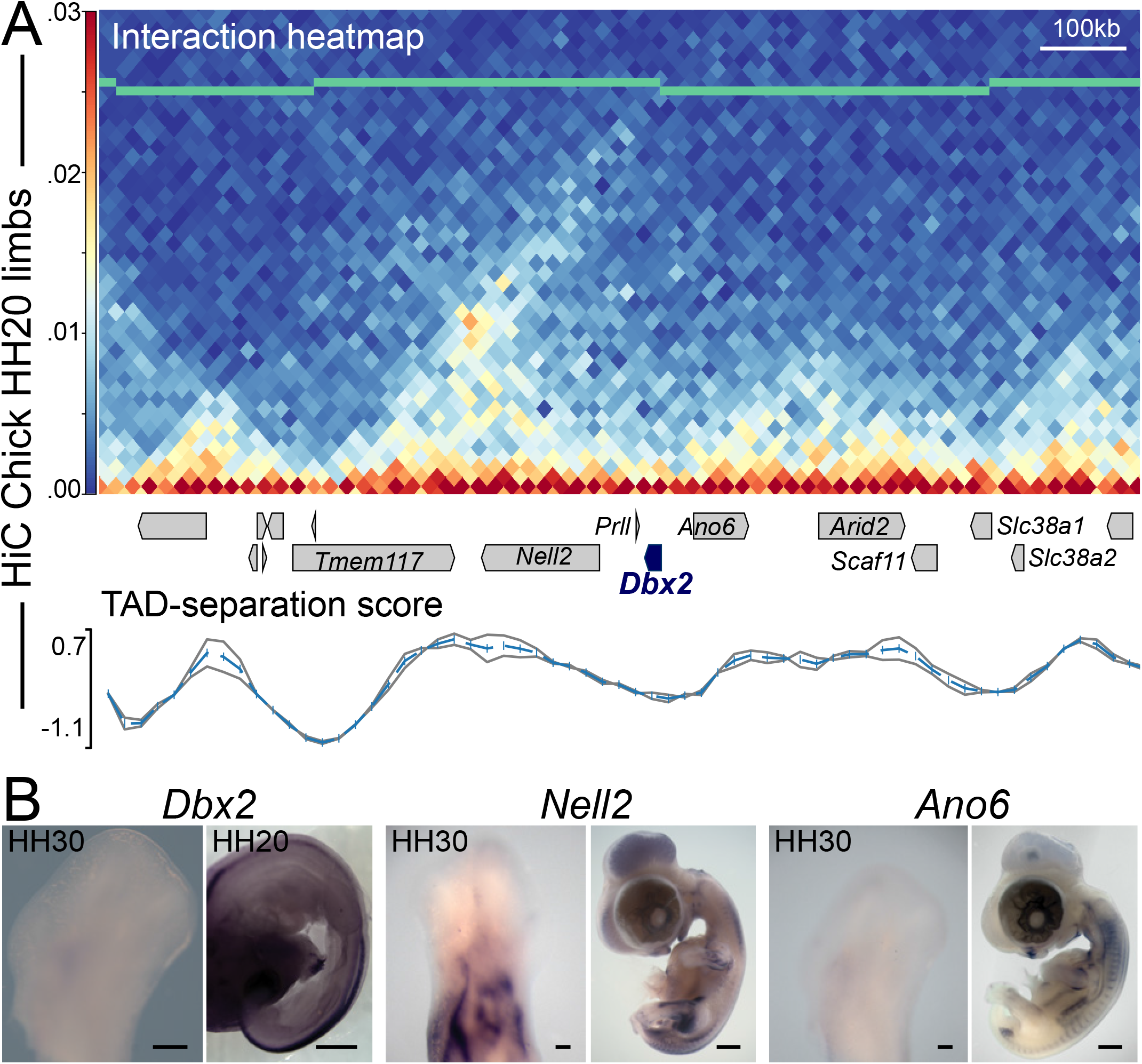
TAD structure and expression of the chicken *Nell2*/*Dbx2*/*Ano6* genes. **A**. Hi-C map of the *Dbx2* genomic region in chicken HH20 wing buds and graphs showing the TAD-separation score (bottom) using standard (gray lines average in blue in the top graph) or a fixed window size of 120Kb (blue line in the bottom graph). Protein-coding gene loci are represented by blue (*Dbx2*) or gray boxes for all other genes. Data from ^58^. TADs called by hicFindTADs are depicted with black boxes. **B**. WISH analysis of *Dbx2, Nell2* and *Ano6* expression in chicken wings and in HH20/HH30 embryos. Scale bar 250µm (limbs)/ 2mm (embryos).

Therefore, we asked whether *Dbx2, Nell2*, and *Ano6* were expressed in the developing wings of chick embryos. We did not observe the expression of either *Dbx2, Nell2* or *Ano6* in the distal domain of embryonic chick limb buds (Fig. 7B, C) by WISH. Although weak expression levels of *Dbx2, Ano6* and *Nell2* transcripts were detected in RNAseq data, they were not specifically increased in the chicken autopod ^58^, in agreement with the idea that digit-specific expression of these genes was acquired after the emergence of the eutherian lineage. Instead, *Dbx2* was expressed in the developing chicken neural tube (Fig. 7C), in agreement with its expression in the mouse CNS and with the presence of the evolutionary conserved, neural tube-specific, mm1571 regulatory element located within the second *Dbx2* intron (Fig. 3A, B; Fig. S3). Likewise, *Nell2* was expressed in the neural tube and somites of both species, in agreement with the function of this gene in sensory and motor neurons differentiation ^59,60^. *Ano6* transcripts were also detected in the paraxial and lateral mesoderm of both species. Therefore, *Dbx2, Nell2*, and *Ano6* expression in embryonic structures others than the developing digits is common to different tetrapod lineages, yet it is associated with different populations of neural and mesodermal precursors, suggesting that their transcription in these structures likely relies on gene-specific regulatory elements (Fig. 7B, C; Fig. S3). These results suggest an evolutionary scenario whereby the acquisition of distal limb enhancers within an ancestral TAD organization led to the co-option of *Nell2, Dbx2* and *Ano6*, in the developing mouse digits. The functional consequences of this co-option remain to be established.

## DISCUSSION

### HOX13 mediated activation of *Dbx2* in digits and its function in distal limb development

In this study, we show that *Hoxa13* and posterior *Hoxd* genes directly activate *Dbx2* expression by binding to different regulatory elements located either within the *Dbx2* introns or in the 30Kb 5’ to *Dbx2*. This is supported by the expression of these former genes in the autopod anlage, which precedes that of *Dbx2* by approximately 24h. Because HOX13 proteins have been proposed to act as pioneer factors ^24^ (see also ^60^), their binding at the *Dbx2* locus may facilitate the access to other transcription factors, thus explaining why some *Dbx2* expressing cells do not express any *Hox13* genes in E13 and E15 distal limbs whereas mice lacking all *Hox13* functions completely loose *Dbx2* expression. Besides binding to the DLE1 to DLE3 sequences, HOXA13 and HOXD13 also bind to various locations within the *Nell2*/*Dbx2*/*Ano6* genomic region. Many such sequences are only partially conserved across the eutherian lineage, in contrast with the high conservation of the DLE regions. While the functional significance of this large HOX13 coverage has not been addressed, it is clearly reminiscent of what was described for the TAD flanking the *HoxD* cluster, suggesting that HOX13 proteins may global regulate the TAD activities at the *Dbx2* locus ^23^.

Mice lacking *Dbx2* function did not show any major skeletal anomaly, neither in the number of phalanges, their length or their ossification pattern, nor in their joints. Also, we did not observe any major limb alterations in the offspring of mice carrying a deletion of the DLE1 sequence, thus suggesting that *Dbx2* is likely not a major mediator of HOX13 function during distal development. The observed embryonic/perinatal lethality of *Dbx2*^*-/-*^ mice could result from defects in the specification of neuronal type in the CNS. However, *Dbx2*^-/-^ mice could display as yet undetected anomalies in the development and/or function of digital tendons and/or ligaments, as suggested by the expression of this gene in precursors identified by the presence of transcripts from *Scx*, a known marker of tendon and ligament progenitor differentiation ^42,62^. Therefore, our results do not support the possibility that the loss of function of *DBX2* alone leads to the hand/foot defects observed in humans carrying a heterozygote deletion of the *NELL2*/*DBX2*/*ANO6* genomic region ^36^. However, the observation that these genes are expressed during embryonic limb development indicates that their combined loss may generate these severe limb alterations. Accordingly, *Ano6* inactivation in mice was reported to affect bone formation and to result in micromelia ^63^. In this context, the loss of the shared DLE1 regulatory element in ungulates may illustrate the flexibility of distal limb structures and their spectacular morphological diversification. Accordingly, we cannot rule out the existence of a human-specific function of the *DBX2*/*NELL2*/*ANO6* genes in hand/foot development.

### Contribution of chromatin architecture and enhancer activity in *Nell2, Dbx2*, and *Ano6* coregulation in developing digits

We used a comprehensive set of scRNA-seq, ChIP-seq and Hi-C data to identify regulatory elements controlling *Dbx2* expression in digits and two such elements (DLE1 and 2) displayed enhancer activity in transgenic mice. The deletion of DLE1 leads to a strong downregulation of *Dbx2* transcripts. The DLE1-3 sequences are evolutionarily conserved across eutherians, while they were not identified in other vertebrate species. Together with our observation that *Dbx2* is not expressed in developing embryonic chicken extremities, this suggests that limb-specific *Dbx2* expression evolved in the mammalian lineage. Also, the comparison of Hi-C interaction profiles at the *Dbx2* loci between mouse and chick revealed a similar TAD organization (Fig. 8B), which likely originated early in the tetrapod lineage, although we cannot exclude that more subtle changes in TAD architecture also contributed to the evolution of *Nell2*/*Dbx2* and *Ano6* expression in digits. However, while in chick the *Dbx2, Nell2* and *Ano6* promoters are regulated mostly by locus-specific short or mid-range regulatory interactions, the mouse DLE can co-regulate these three genes despite their location in different TADs.

**Figure 8.**
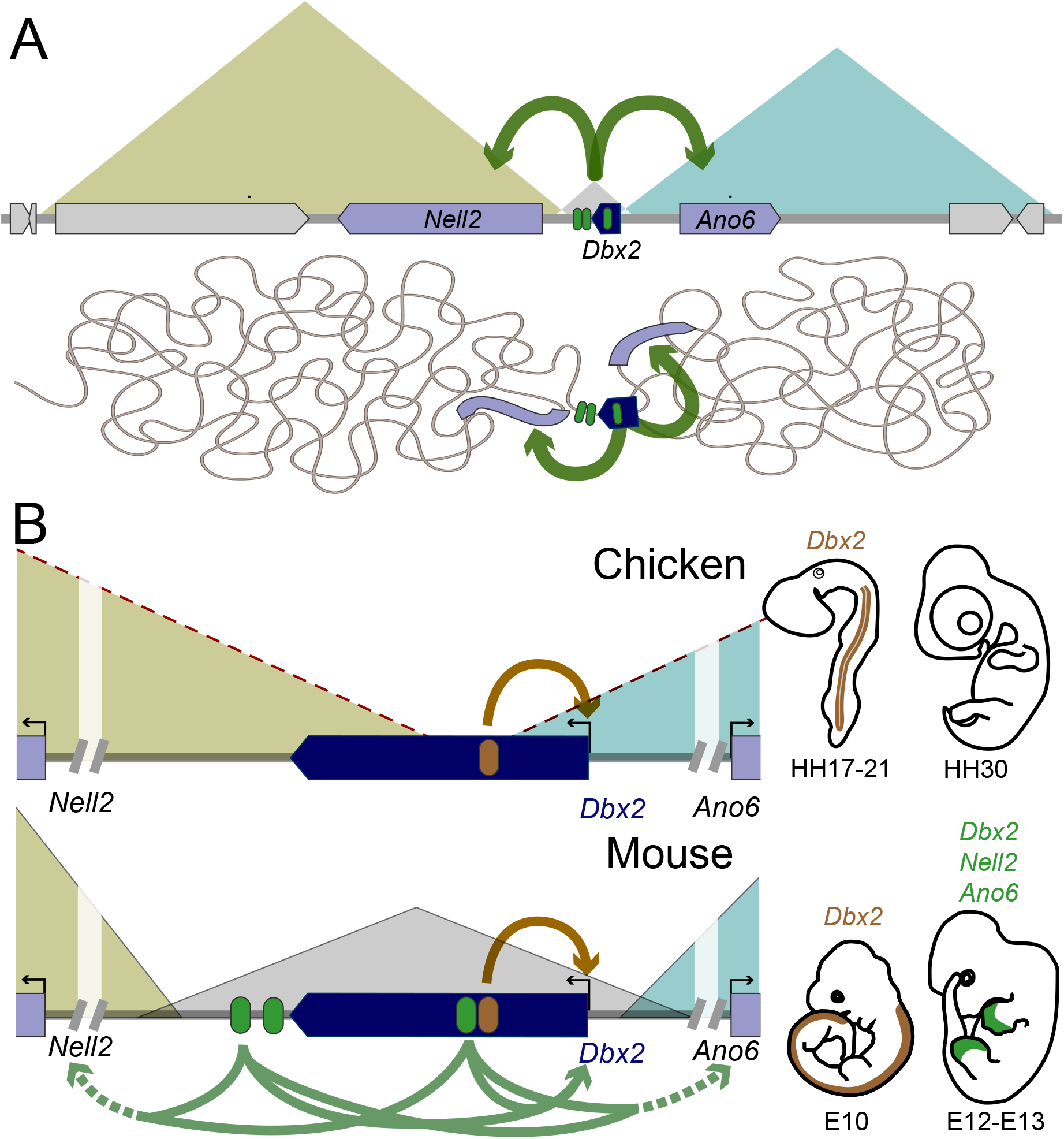
*Dbx2* regulation in mouse and chicken. **A**. Scheme depicting the TAD architecture of the mouse *Dbx2* genomic region (top) and its 3D organization (bottom). The *Dbx2* and *Nell2*/*Ano6* loci are depicted by blue and light purple boxes, respectively. All other genes are in gray boxes. DLE1-3 elements are depicted in green. Green arrows indicate the regulation of DLE1-3 over *Nell2* and *Ano6* genes. **B**. Schemes depicting the TAD organization of the mouse and chicken *Dbx2* genomic region and its regulation. DLE1*-*3 and Vista mm1571 (or its chicken orthologous) elements are depicted by green and brown round boxes, respectively. Because of the low resolution of the chicken Hi-C, it was not possible to precisely resolve the location and extension of the *Dbx2* interTAD domain (approximate TADs limits are depicted by red dashed lines). Green and brown arrows depict the DLE1-3 and neural tube enhancer regulatory activity, respectively. No expression of *Dbx2, Nell2* or *Ano6* is scored in the distal limb/ wing of the chick embryo. In the mouse, the DLE1-3 regulate the expression of *Dbx2, Nell2* and *Ano6* in the developing distal limb.

This organization is reminiscent of that observed at the *HoxD* cluster, yet with an inversion of functionalities. At the *HoxD* locus, two flanking TADs contain distinct enhancers, which act in an exclusive manner upon *Hoxd* genes located at the TAD boundary (e.g.^64–66^). At the *Dbx2* locus, the regulatory elements are located at the TAD boundary and can interact with target genes located within the two adjacent TADs, thus providing a first example of such a regulatory architecture. However, the functional contribution of this organization, as well as the mechanisms whereby the DLE sequences can differentially interact with the *Nell2*/ *Dbx2* and *Ano6* promoters, remain to be determined. Recent reports have used chromosome engineering to analyze the insulating effect of TAD boundary regions ^67–69^, supporting the conclusion that TADs are domains where enhancer-promoters contacts are favored, if not constrained. Our results suggest that, in some cases, enhancers located in between TADs may be selected to interact with either TAD depending on the context. Accordingly, it was recently shown that boundary elements can play an important role in allowing the establishment of interTAD promoter-enhancer interactions in drosophila embryos ^70^, yet with a mechanism substantially different from the one proposed here. Another non-exclusive possibility is that *Dbx2* would be expressed in developing digits as a bystander effect due to the activity of the neighboring limb enhancers^71^.

## EXPERIMENTAL PROCEDURES

### Mouse strains and transgenesis essays

Mice were kept and handled following good laboratory practices. Mutant strains were maintained in heterozygosis. The *Hoxd13, Hoxa13*, Del(1-13), De(8-13), and Del(9-12) mutant mouse lines (Fig. 4) were previously described ^17,20,72–74^.

To generate *Dbx2*^+/-^ and *DLE1*^+/-^ mutant lines pairs of specific sgRNA targeting both sides of the *Dbx2* third exon (CTGCTGTTGAAAGTAGGACT; CCACTGTTCTGAGAGTCCGA) and the *DLE1* enhancer (GAAAAGGAAGACCACCCGTG; AGGGGCTAGAGATCTCCCAG) were co-electroporated, together with the Cas9 protein (TruecutV2; Thermofisher), in fertilized mouse oocytes. To screen for each mutant allele, we designed specific primer pair flanking the *Dbx2* third exon and enhancer (DLE1_F: ACACACAGATAAATGCACGTGAAGTG; *DLE1*_R: GGAGGGCCACTCTTAGGTGTG). In each case, we selected F0 mouse mutant carrying a deletion of 377bp (chr15: 95632232-95632608, mm10) spanning the whole *Dbx2* third exon, and a 1015bp deletion (chr15:95600674-95602176, mm10) encompassing the *DLE1* sequence. Mutants mice were backcrossed with Wt B6CBAF1 mice. Mutant F1 and F2 mice were selected using specific genotyping primers for their respective Wt and mutated alleles (*Dbx2*_F: GGAACTCCCACCTTCGACTGACTG/ ACTGTTGATTAGGGCTGGGCTTTGA; Wt alleles: 756/ mutant allele 388bp; *DLE1*_Wt: GGAGTGAGGTTGTGCCAAGA/ ACCTGTAAGCCAACCCCTAC; *DLE1*_Mut: ACACACAGATAAATGCACGTGA/ GAGGGCCACTCTTAGGGTGG).

For the transgenesis essays, the Tg*DLE1*::LacZ and Tg*DLE2*::LacZ plasmids were linearized with NotI and KpnI. The fragment encoding the enhancer, b-globin minimal promoter and LacZ reporter was gel purified and injected in the masculine pronucleus of fertilized oocytes. Transgene injections were performed by the transgenesis platform of the University of Geneva. F0 embryos were dissected at E12-E13 and stained for LacZ activity.

### Probe, transgene and sgRNA cloning

The sgRNA targeting guides were generated by annealing complementary pairs of oligonucleotides and cloned into the pX330 vector as described in ^65^.

The plasmid encoding for the mouse *Dbx2* RNA probe was a gift from Thomas Jessell (Addgene plasmid 16288; ^33^). Instead, specific primers were used to amplify a portion of the transcribed region of the mouse *Nell2, Ano6, Gdf5* and *Mkx* genes as well as of the chicken *Dbx2, Nell2* and *Ano6* orthologs. In each case, the PCR products were cloned into the pGEM-Teasy plasmid and sanger sequenced. For the chicken *Dbx2, Nell2* and *Ano6* genes, primers were designed based on the exon/intron structure predicted from the UCSC Non-Chicken Refseq genes, spliced EST, Chicken mRNA and Ensembl gene prediction.

For the transgenesis assays, the DLE1 /DLE2 sequences were amplified with specific primers (*DLE1* Fw: ATCCTGCTGTCTCTGGCTTTCAT/ GGGATCTGATGCATGTAGTGGAATTC; *DLE2* Fw: TCCAAGTTCTGTCTTCTAGGGCA/ GGATTGTGTATTAACCAGGACCGA) and cloned into the pSK-bglob::LacZ reporter plasmid ^23^, generating the Tg*DLE1*::Lacz and Tg*DLE2*::LacZ reporter vectors.

### Probe and sgRNA preparation

For the sgRNA transcription, we PCR amplified the sgRNA sequence cloned into the px330 plasmid using a T7 promoter containing primer and a universal reverse oligonucleotide (TAATACGACTCACTATAG). PCR products were gel purified and transcribed in vitro using the HiScribe™ T7 High Yield RNA Synthesis Kit (NEB). The transcribed sgRNAs were purified using the RNeasyTM mini kit (Qiagen).

Specific probes for the different genes analyzed were synthesized in vitro by linearizing the respective coding plasmids using specific restriction enzymes and by in vitro transcribed with either T7/T3 or Sp6 RNA polymerase. The probes were purified using the RNA easy mini kit.

### Gene expression analysis

For the qPCR analysis, pairs of E10/E11/E12/E13 mouse DFLs, as well as HH30-31 chicken distal wings, were microdissected and stored in RNAlater. Toral RNA was extracted from each pair of DFLs/ distal wings using the Qiagen microRNA extraction kit and retro-transcribed using the Promega GOscript reaction mix with random primers. Gene expression levels were measured by real-time qPCR using the SYBR® Select Master Mix for CFX (Thermofisher), and specific primer pairs for the mouse *Dbx2, Hoxa13, Hoxd13* genes as well as for the chicken orthologs *Hoxd13* and *Dbx2* (Table I). The mouse *Hmbs* and chicken *Gapdh* housekeeping genes were used as internal controls for the normalization of gene expression levels (2^-(ΔCt)^). WISH experiments were performed as described in ^75^.

### Skeletal preparation

Alcian blue and alizarin staining was performed as described in ^54^. Briefly, P7 mouse pups cadavers were eviscerated and skin and fat tissues were removed as much as possible. After 48h fixation in ethanol, cadavers were stained alcian blue solution (150 mg/l alcian blue 8 GX in 80% ethanol and 20% acetic acid) for two days and washed in 100% ethanol overnight. Subsequently, they were cleared for at least 3h in 2% KOH solution and stained for 2h in 50 mg/l alizarin red / 2% KOH solution. Finally, they were washed in 2% and 1% KOH solution and progressively dehydrated to 100% glycerol solution.

### Hi-C/ ChIP seq/ scRNA-seq data analysis 4C-seq interaction profiling

All scripts used to analyze data and generate figures are available at https://github.com/lldelisle/scriptsForBeccariEtAl2021. The calculations were performed using the facilities of the Scientific IT and Application Support Center of EPFL.

For the chicken Hi-C analysis, the raw forelimb and hindlimb data were extracted from GEO (see Table II) and processed independently using HiCUP v0.8.0 ^76^ on galGal6. Valid pairs were obtained using a custom python script. Both valid pair files were merged before analysis. Valid pairs from Hi-C carried out on mouse material were downloaded from GEO (see Table II).

Valid pairs of each study were loaded in a cool file using cooler version 0.8.10 ^77^ using a resolution of 5Kb, 20Kb or 40Kb. The TAD-separation score and the domains were obtained using hicFindTADs version 3.5.2 ^78–80^ with --minBoundaryDistance 100000 and either default parameters for the choice of window size or a fixed window size of 120Kb. Plots were obtained using pyGenomeTracks version 3.6^78,81^.

ChIP-seq paired-end (PE) fastq of HOXA13 and HOXD13 data, as well as single-read (SR) fastq from H3K27me3 and H3K27ac and corresponding inputs were downloaded from GEO (see Table II). Adapter sequences and bad quality bases were removed with Cutadapt^82^ version 1.16 with options -a GATCGGAAGAGCACACGTCTGAACTCCAGTCAC -A GATCGGAAGAGCGTCGTGTAGGGAAAGAGTGTAGATCTCGGTGGTCGCCGTATCATT -q 30 - m 15 (-A being used only in PE datasets). Reads were mapped with bowtie 2.3.5^83^ with default parameters on mm10. Alignments with a mapping quality below 30 were discarded with samtools view version 1.9^84,85^. For HOXA13 and HOXD13, coverage and peak calling were computed by macs2 version 2.1.1.20160309 with options --call-summits -f BAMPE -B. Coverage was then normalized by the number of million fragments used in macs2 coverage. For histone marks, coverage and peak calling were computed by macs2 with options -f BAM --nomodel --extsize 200 --broad using the BAM of input in -c. The coverage was then normalized by the number of million tags used in macs2 coverage. Plots were obtained using pyGenomeTracks version 3.6^78,81^. For DFL_E12_H3K27ac, the two replicates were averaged.

For the scRNA-seq, matrices with counts were downloaded from GEO (see Table II). UMAP and expression plots were obtained using Seurat package version 3.2.2^86^ on each dataset individually.

We performed our 4C-seq experiments according to ^87^. Briefly, 12 pairs of wildtype DFLs or PFLs were dissected, dissociated with collagenase (Sigma Aldrich/Fluka) and filtered through a 35 micron mesh to isolate single cells. Cells were fixed with 2% formaldehyde (in PBS/10%FBS) for 10 min at room temperature and the reaction was quenched on ice with glycine. Cells were further lysed with 10 mM Tris pH 7.5, 10 mM NaCl, 5 mM MgCl2, 0.1 mM EDTA, 1x Protease inhibitor cocktail to isolate nuclei and stored at -80°C. Nuclei from pools of at least 10 distal or proximal limbs were digested with DpnII (New England Biolabs) and ligated with T4 DNA ligase HC (Promega) in diluted conditions to promote intramolecular ligation. Samples were digested again with NlaIII (New England Biolabs) and ligated with T4 DNA ligase HC (Promega) in diluted conditions.

These templates were amplified using Expand long template (Roche) and inversed PCR primers flanked with adaptors allowing multiplexing (Table III). Barcodes (4bp) were added between the Illumina adaptor and the specific DpnII primers. Libraries were prepared by means of 8–10 independent PCR reactions using 70–100 ng of DNA per reaction. PCR products were pooled and purified using the PCR purification kit (Qiagen). Multiplexed libraries were sequenced on Illumina HiSeq 2500 at the Sequencug platform of the University of Geneva to obtain 100 bp single-end reads. Demultiplexing, mapping and 4C-seq analysis were performed using a local version of the pipeline described in ^88^, on the mouse assembly GRCm38 (mm10). The profiles were smoothened using a window size of 11 fragments and normalized to the mean score in +-5 Mb around the viewpoint. When multiple independent biological replicates were available, average 4C-seq profiles were calculated.

Data are available in GEO (accession number: GSE161386).

## ACKNOWLEDGMENTS

We thank the transgenesis and sequencing platforms of the University of Geneva. We also thank Aurélie Hintermann for help in chick embryo dissection and for discussions.

