## Supplementary figures for "HOX13-MEDIATED *DBX2* REGULATION IN LIMBS SUGGESTS INTER-TAD SHARING OF ENHANCERS"

By Leonardo Beccari et al.

This document contains:

Supplementary figures 1-13,

and associated references.

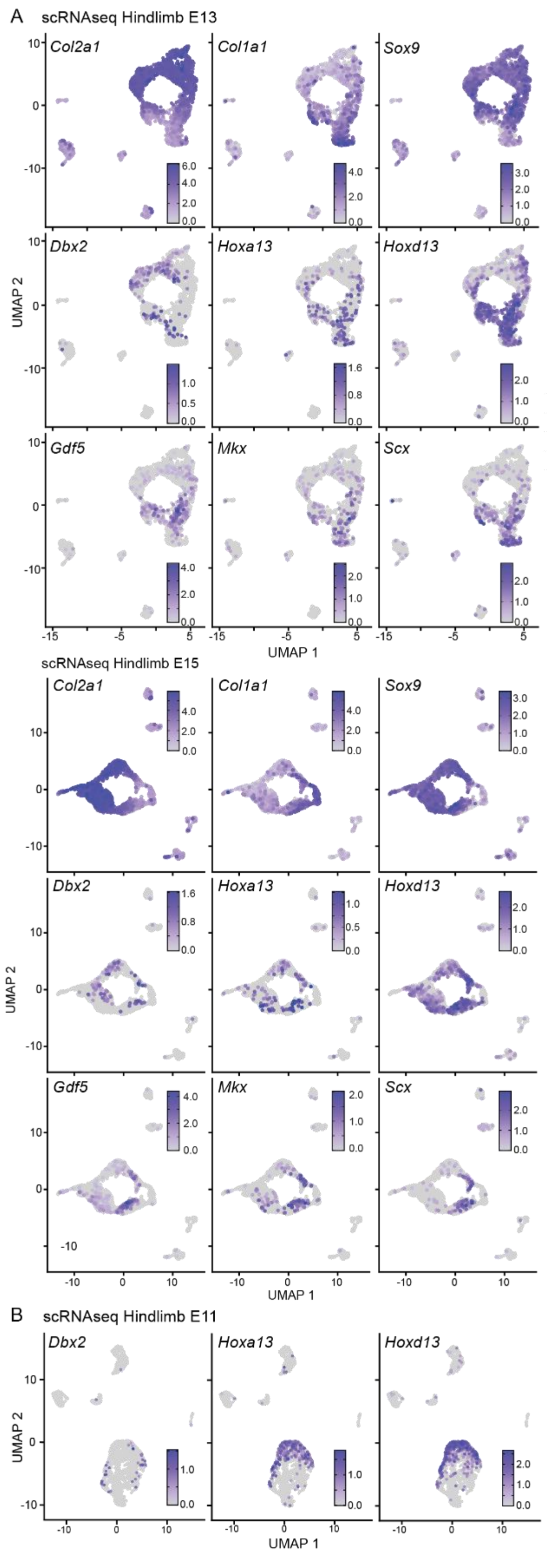


**Supplementary Figure S1. Single-cell RNAseq analysis of *Dbx2*+ cell populations in developing hindlimbs.** UMAP representations of the scRNAseq data from mouse E11 (**A**) E13 (**B**), and E15 (**C**) mouse hindlimbs [1] showing the expression of *Dbx2*, *Hoxa13* and *Hoxd13*, as well as of different joint (*Gdf5*) and tendons/ligaments (*Mkx*, *Scx*) markers [2–4].

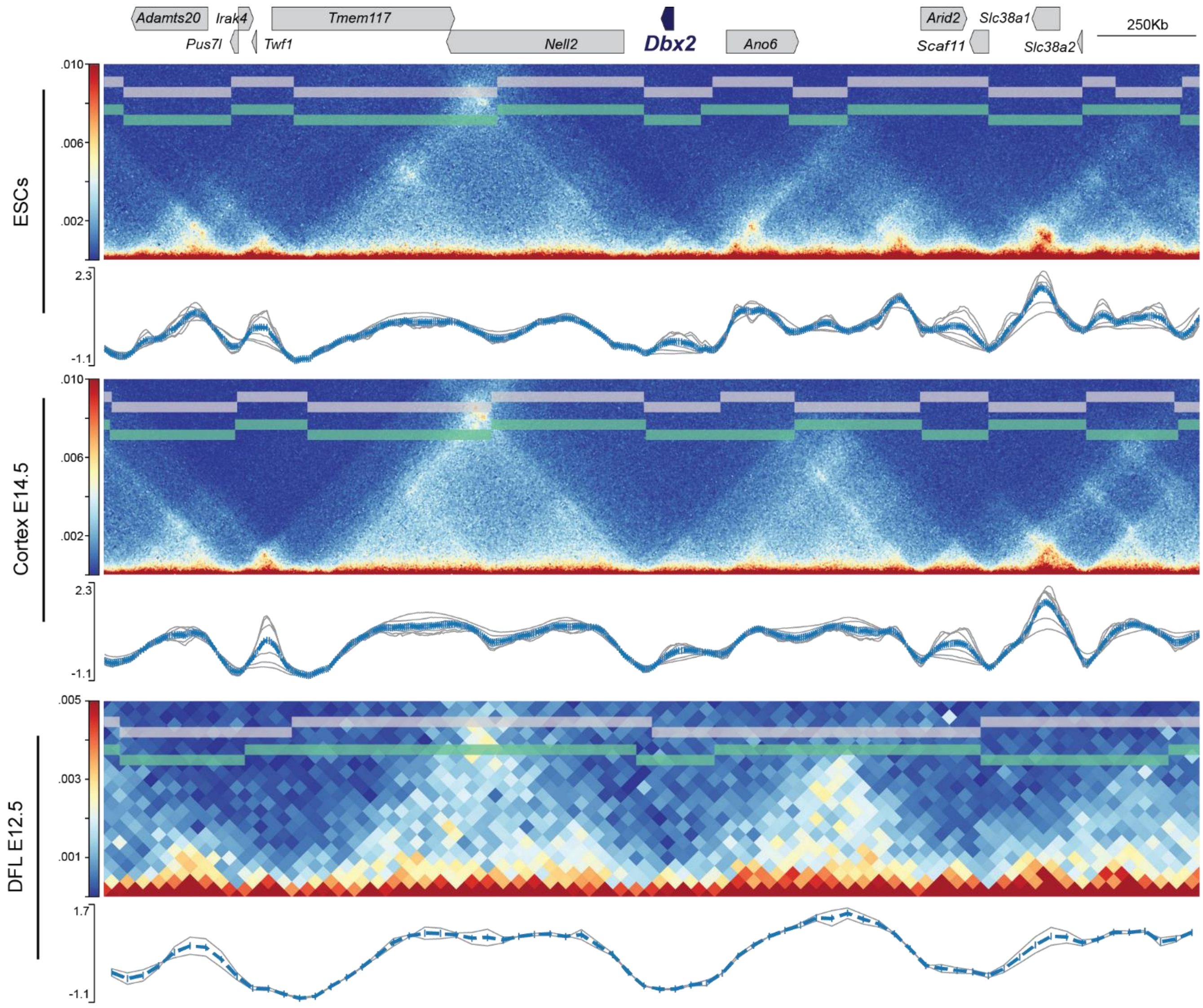


**Figure S2**

**Supplementary Figure S2. TAD organization around the *Dbx2* locus. A.** High resolution (5kb bin size) Hi-C map of the *Dbx2* genomic region in mouse ES cells (top) and E14 embryonic cortex (bottom), and graphs showing the TAD-separation score based on the HicFindTADs algorithm using different window size values (the curve calculated using standard parameters is displayed in gray and the average in blue). Data from [5]. **B.** 40kb resolution (bin size) Hi-C map of the *Dbx2* genomic region in E12 mouse limb buds and graphs showing the TAD-separation score (as in **A**). Data from [6]. On top, the gene loci are represented in blue (*Dbx2*) or gray boxes for other genes. (A,B).

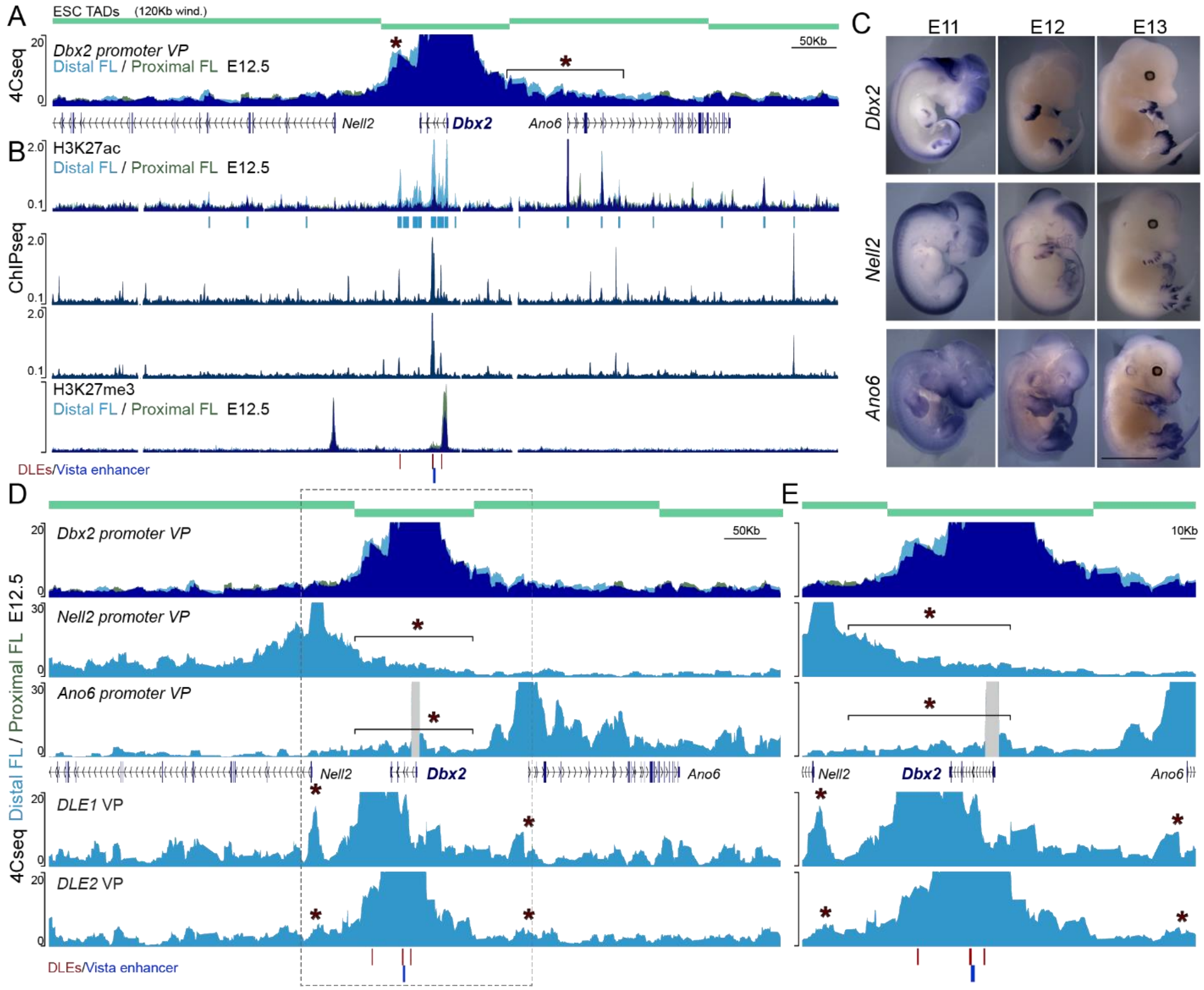


**Supplementary Figure S3. Characterization of *Dbx2* regulation in mouse developing limbs. A.** zoomed-in view of the 4Cseq interaction profiles of the *Dbx2* promoter (from Fig. 2A). Asterisks label the region displaying an increased contact frequency in the distal *versus* proximal limb bud. The TADs (data from [5]) are on top as light blue boxes **B.** ChIPseq analysis of H3K27ac and H3K27me3 marks in distal (light blue) and proximal (green) forelimbs, and HOXA13/HOXD13 binding profiles over the *Dbx2* genomic region. Data from [7,8]. Profile overlap in A and B is in dark blue. **C**. WISH analysis of *Dbx2*, *Nell2* and *Ano6* expression at different developmental stages. **D, E**. Zoomed-in views of the 4Cseq profiles showing the interactions of the *Dbx2*, *Ano6* and *Dbx2* promoters, as well as of the *DLE1* and *DLE2* sequences (see Fig, 6D) in distal (light blue) and proximal (green) forelimb buds (profile overlap is in dark blue). DLE1-3 elements are depicted with red boxes. The vista mm1571 element is in blue. Probably artefactual PCR product is depicted in gray. The region displayed in E is marked by a dashed rectangle in D.
